# The impact of AlphaFold on experimental structure solution

**DOI:** 10.1101/2022.04.07.487522

**Authors:** Maximilian Edich, David C. Briggs, Oliver Kippes, Yunyun Gao, Andrea Thorn

## Abstract

AlphaFold2 is a machine-learning based program that predicts a protein structure based on the amino acid sequence. In this article, we report on the current usages of this new tool and give examples from our work in the Coronavirus Structural Task Force. With its unprecedented accuracy, it can be utilized for the design of expression constructs, de novo protein design and the interpretation of Cryo-EM data with an atomic model. However, these methods are limited by their training data and are of limited use to predict conformational variability and fold flexibility; they also lack co-factors, posttranslational modifications and multimeric complexes with oligonucleotides. They also are not always perfect in terms of chemical geometry. Nevertheless, machine learning based fold prediction are a game changer for structural bioinformatics and experimentalists alike, with exciting developments ahead.

## Introduction

### Cryo-EM structures are an under-determined problem with noisy data

In recent years, electron cryo microscopy has opened up possibilities to see and understand many large molecular machines and membrane complexes which were previously inaccessible to the structural biology community^1^. Singleparticle electron cryo microscopy (Cryo-EM) does not require crystallization, uses very small amounts of material and is applicable to a wide range of macromolecule sizes. It also permits us to study fibrils, membrane proteins and viral assemblies, structures which are typically inaccessible by crystallography. The so-called “resolution revolution”^2^ (Fig.1) led to the 2017 Nobel prize for developing cryo-electron microscopy as researchers overcame limitations in sample preparation, detector technology and image processing^3^. Very recently, the possibility to determine atomic resolution structures has been experimentally demonstrated^4–6^, a very exciting and long predicted development. However, solving an atomic structure from Cryo-EM micrographs remains an underdetermined problem with noisy data: molecular particles are hard to make out with the bare eye in the micrographs as the signal is so weak, and the inherent flexibility of molecules as well as thousands of atomic positions in each particle represent a huge parameter space for the atomic models we utilize for interpretation. For very large structures, computation can be an obstacle. Furthermore, for many structures, only low-resolution data are available, as inherent flexibility and particle heterogeneity blur reconstruction maps. Another kind of revolution may now enable us to obtain more from such low-resolution data and has many implications for Cryo-EM, as it allows us to add prior knowledge in an unprecedented way to the experimental data:

**Fig. 1.**
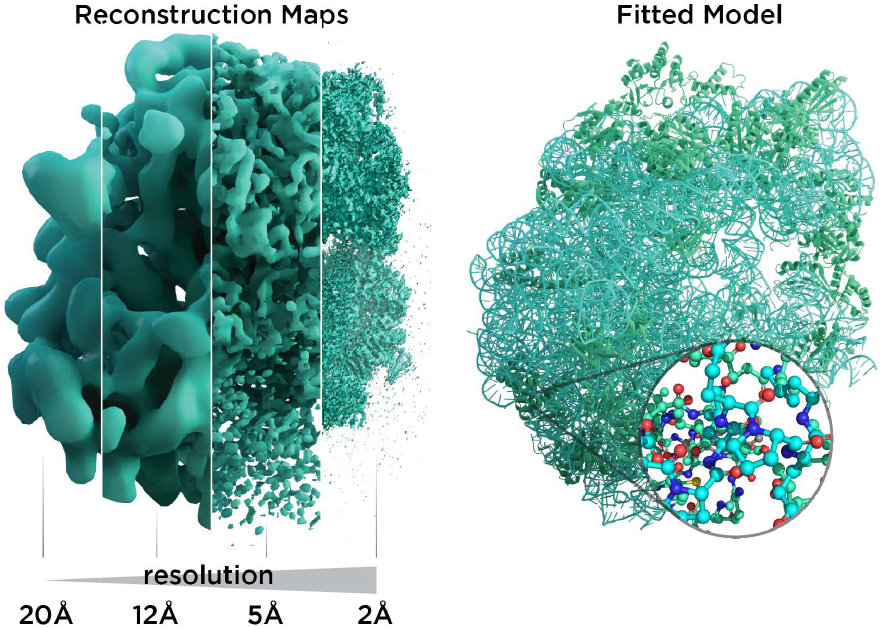
Resolution revolution in Cryo-EM. Left: Reconstruction maps shown for the 50S ribosome with increasing level of detail from left to right as the field evolved. Right: A typical atomic model used for interpretation of reconstruction maps. Picture courtesy of Andrea Thorn and Thomas Splettstößer.

### AlphaFold

AlphaFold^7^ is a machine-learning (ML) based program that predicts a protein structure based on the amino acid sequence. Its second generation, AlphaFold2‡ won the CASP 14 challenge in 2020 in predicting protein folds, outperforming all previous algorithms for fold prediction from individual protein sequences. Chemical and Engineering News wrote “At this year’s competition, two-thirds of the protein structures predicted by AlphaFold were within experimental error. Basically, these structures were as good as the ones researchers could obtain through their laboratory techniques.”^8^

These experimental techniques, such as Cryo-EM, are currently the main method for structure solution, and will certainly remain in use, in particular in studies focussing on large macromolecular assemblies with complex interactions. However, some structure determinations can now be replaced by ML-based fold prediction. AlphaFold is already used to solve, model and improve protein structures^9–12^, and *ab initio* model building in electron density or Cryo-EM reconstruction maps will no longer be a major challenge for experimental structural biology. This will have significant impact, not only for singleparticle electron cryo microscopy and crystallography, but also on NMR, electron tomography and small angle X-ray scattering. Most developments detailed here are very recent and ongoing, with many references being not yet peer-reviewed.

Therefore, selected examples from our work as part of the Coronavirus Structural Task Force will be used to highlight major points. This task force is an initiative of 26 structural biologists and students who focus on evaluating, improving and consolidating our knowledge of SARS-CoV-2 by critically assessing and disseminating structural information about SARS-CoV-2 proteins^13^, which occasionally involves machine learning based fold prediction as well.

## Applications in experimental structural biology

Machine learning-based fold prediction is an extremely valuable tool for structural biologists. We will highlight how it can be used in connection with experimental work, but also which limitations apply. AlphaFold estimates per-residue confidence on a scale from 0 – 100, with higher values being better; this score is called *predicted local distance difference test* (pLDDT). It is stored in the B-factor fields of the model file. The AlphaFold pipeline begins with the generation of a deep multiple sequence alignment and a matrix of distance restraints derived from covariant evolutionary differences in the amino acid sequences. These inputs are then passed through a neural network block called “Evoformer” that produces processed multi sequence alignments and an array of pair-pair distances. The following structure block uses these inputs to position and refine the relative location of each amino acid in 3D space. This whole process is then recycled several times to produce the final model.

### Construct design

Perhaps the first application that comes to mind for usage of AlphaFold for experiments is design of expression constructs. Both the actual geometry of a predicted fold as well as the pLDDT for each residue can be utilized to get an idea about the less compact or ordered regions and the more ordered, or folded, parts of a given protein sequence, where backbone hydrogen bonds are saturated. Often, omitting less ordered regions from a protein sequence is beneficial to design well behaved recombinant proteins for structural study. In order to ensure solubility, ML-based fold prediction may also give information on surface residues of a given construct, and can permit biologists to better decide where a domain starts and ends in the sequence.

One example for this is the *betacoronavirus-specific* marker domain of the protein nsp3. The roughly 200 residues long domain had previously been predicted to be mostly disordered^14^. However, an AlphaFold prediction revealed a stable folded region of ~80 residues surrounded by large disordered linkers, and recently, this region was experimentally confirmed to be folded (PDB ID 7T9W).

### *De novo* protein design

*De novo* protein design has been a goal of the structural biology community for decades, but was limited by the need to experimentally validate each and every new sequence in order to test hypotheses. With more reliable fold prediction available, this has now changed^15,16^. However, at the time of writing this article, no experimentally validated folds designed by AlphaFold have yet been published.

An obvious translational outlet from *de novo* protein design would be the creation of “biologics”, such as antibodies^17^ or DARPins^18^. To date, creation of such molecules has favoured time- and resource-intensive experimental techniques such as ribosome-scanning^19^. If design of biologic therapeutics could be carried out computationally, this would represent a huge advance for potential therapeutics but also research and diagnostic tools.

Another related use of protein design technology would be vaccinology. At the beginning of the COVID-19 pandemic, Pfizer/BioNTech and Moderna vaccines included Spike-protein stabilising mutations to increase the half-life and thus the efficacy of their vaccines^20^. Mutations to SARS-CoV-2 Spikeprotein were introduced based upon knowledge of structures of the related SARS-CoV-1 and MERS Spike proteins, but it is clear that *de novo* design of stabilised immunogens would be beneficial to future vaccine-design work.

### Structure Solution

ML-based fold prediction can be used to solve structures in crystallography by molecular replacement,^11^ and to fit electron density and Cryo-EM reconstruction maps. Terwilliger *et al.* have designed an iterative AlphaFold modelling pipeline that iterates between AlphaFold modelling and refinement against experimental data (either Cryo-EM or macromolecular crystallography) to yield improved models^12^. Machine learning based fold prediction has not been used for validation yet, but it is being used to actively improve and correct experimental structure determination^9–11,21^.

## Limitations of machine learning based fold predictions

### Impact of Training Data

AlphaFold is a supervised machine learning method, hence its neural networks are trained with data for which the fold is known, allowing its constituent neural networks to learn how to derive the fold from the sequence. The correct choice of training data is crucial to any machine learning project^22^. In the case of AlphaFold, the training data are from the world-wide Protein Data Bank (PDB)^23^, supplemented with selected “self-training” predictions^24^ the source of which remains somewhat vague^7^. However, the PDB data have some very specific properties which affect the performance of AlphaFold, mainly the lack of intrinsically disordered structures and a very limited sampling of conformational space^25^.

#### Dark Proteome

The “dark proteome” is comprised of proteins with no stable fold that represents a well-defined threedimensional structure, an estimated 44% of proteins in eukaryotes and viruses^26,27^. These are predominantly intrinsically disordered proteins which are thought to contribute to defence and signalling, sometimes becoming ordered when interacting with other macromolecules. As we have very little data about these proteins in a structural sense, besides their sequence, their structures as well as function cannot (yet) be modelled by machine learning. However, to an extent AlphaFold is able to predict where disorder occurs, as the pLDDT in such regions will be low^28^. Williams et al.^29^ differentiate structures which AlphaFold cannot predict well into those which still look “like” protein and so-called “barbed wire” folds where residues are just lined up next to each other in a nonsensical conformation with little hydrogen bonding. They speculate that in the latter case, there was little evolutionary covariance in that region.

One exciting example of such a protein is the SARS-CoV-2 nucleocapsid, the complete structure of which is still unknown, but which is essential for the viral infection cycle, and hence an important drug target against COVID-19^30^. The nucleocapsid protein has two ordered domains (which were experimentally determined^31^) and three intrinsically disordered regions: the N-terminal (residues 1-47), C-terminal (365-419) and Linker (175-247) domains. These regions may order upon binding of RNA, which nucleocapsid protects as part of the virion^32^. Nucleocapsid has also been implicated in RNA regulation. AlphaFold simulations of the entire sequence are a very typical example of such a prediction, where ordered compact domains are clearly separated from low confidence disordered areas which show mostly “barbed wire” - with the exception of a leucine-rich helix inside the conserved flexible linker (Fig. 2) and a lower confidence helix in the C terminus. Interestingly, the predicted helix in the Linker domain is the site of the G215C mutation in the delta variant of SARS-CoV-2. This mutation is likely to stabilize conserved transient helices^33^. If machine learning based fold prediction could give us more insight about what happens when RNA is bound, this may shed light on nucleocapsid’s role in the cycle of infection.

**Fig. 2.**
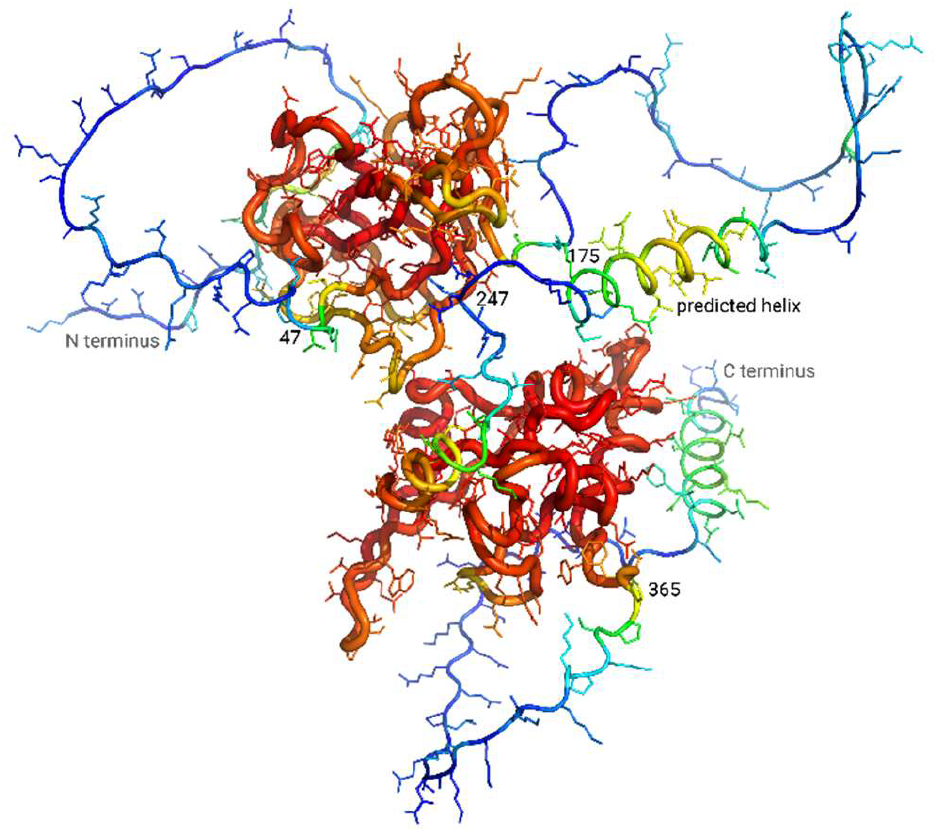
AlphaFold prediction for SARS-CoV-2 nucleocapsid. High pLDDT in red, low pLDDT in blue. While the two known domains are predicted with a high confidence (red), the N-terminus, C-terminus and linker domain are showing a so-called “barbed wire” structure where AlphaFold could not find a solution and has low confidence (blue), with the exception of two helices in the C-terminus and the linker domain. Picture courtesy of Lisa Schmidt and Andrea Thorn.

#### Sampling conformational space

Many macromolecules exist in multiple conformations and are required to adopt these conformations as part of their biological function, particularly enzymes and membrane transporters.

Even when the structural biologists report only one structure from a Cryo-EM experiment there is a wealth of hidden biological information in the Cryo-EM dataset, which can, in principle, be used to observe structural changes and determine free-energy landscapes^34^. Currently, if structures of different states of a molecular machine are needed, the usual approach is to collect and process enough data to determine individual high-resolution reconstruction maps for each state, which are then interpreted individually^35^. However, the fact that many molecular movements are more of a continuous process than jumps between discrete states can make this classical approach problematic^36^.

Indeed, the PDB arguably contains mostly stable conformations, as the majority of structures are derived from vitrified particles (in the case of Cryo-EM) or readily crystallizable proteins (in the case of crystallography). As AlphaFold was trained on this inherently biased training data, it could be assumed that, as a consequence, it can tell us little about different conformations and the inherent flexibility or movement of proteins. There are occasional exceptions, for example, when there is a homomultimer as opposed to a monomer^37^ (see section on “complexes with other proteins”, below). A recent study by del Alamo and colleagues^38^ showed that by reducing the depth of the multiple sequence alignment and limiting recycling of models within the AlphaFold pipeline, they were able to produce multiple fold predictions that appeared to bridge experimentally determined conformations of the same protein, demonstrating that this technique samples conformational space for a range of G-protein coupled receptors and transmembrane transport proteins. As both these classes of protein are important drug targets^39^, this is an important step towards leveraging AlphaFold for translational applications.

It would be highly desirable to get a better grip on the conformational variability information which is present in crystallographic and Cryo-EM data, but not currently really meaningfully used^36,40^ and then to utilize these data to enhance training data for neural networks like AlphaFold.

### Effects of mutations

It is still being debated whether AlphaFold is suitable^41,42^ or not^43,44^ to predict the outcomes of point mutations, with the EBI advising on its homepage that the latter is the case. A preprint by Zhang and co-workers^41^ suggests that, whilst there is no correlation between pLDDT and the destabilising effect of a point mutation, there is however a correlation when looking at ΔΔG or ΔΔTM values derived from the AlphaFold models, which would allow prediction of the destabilising effects of point mutations on protein structure.

### Ideal geometry

Currently, AlphaFold does not produce geometrically perfect structures, even if there are good reference structures in the PDB. SARS-CoV-2 main protease is arguably the most structurally researched drug target for COVID-19, with 485 experimentally determined structures deposited in the PDB. However, when comparing the PDB entry 6YB7 for main protease with the AlphaFold prediction of the same sequence, AlphaFold produces four additional Ramachandran outliers and a higher number of atoms that come too close to each other (with a Clashscore of 16 instead of 2.5 for the experimentally measured and manually built structure, respectively). The predicted fold also has 88 bad bonds and angles as opposed to just 2 in the PDB entry. Some of these occur in the C-terminal region which has a low pLDDT, but they also happen in regions with higher confidence values. It is yet unclear if the deviations from ideal geometry produced by AlphaFold differ from those a human would introduce when modelling a fold to a reconstruction map, and how representative such results are. It might be interesting to investigate this, for example by training a so-called adversarial network^45^ that differentiates between AlphaFold and human-made structures.

### Complexes with other proteins

AlphaFold multimer^46^ is a modification of Alphafold to allow modelling of protein oligomers. The adaptations include pairing the multi-sequence alignments either over n-copies of the monomer for homo-oligomeric cases, or all sequences for the hetero-oligomeric cases, and considering permutation symmetry of homo-oligomeric sequences. Currently this methodology appears to have more success with homooligomers than hetero-oligomers, and the authors assume that this is because the alignment of the homo-oligomer contains more evolutionary information about the homo-typic interfaces than for hetero-typic interfaces^46^. The authors of Alphafold-multimer also note that it is not yet capable of accurately predicting antibody epitopes^46^. Further development of this technology is highly desirable of course, as there would be considerable synergy with sub tomogram averaging and singleparticle Cryo-EM structure determinations of large multimeric complexes.

### Complexes with ligands or oligonucleotides

AlphaFold is not designed to predict the interactions between proteins and other molecules such as RNA, DNA or smaller ligands and co-factors, or of posttranslational modifications, glycosides chief among them^47^. As a consequence, single-chain prediction may or may not correspond to the structure adopted in a complex, be it ligand-induced or through interactions with another macromolecule. Some fold predictions have “holes” where there should be a co-factor or a coordinated metal ion^48^. Efforts to overcome this limitations are being made in the form of adding ligands geometrically to predicted structures^49^ and neural network based methods may soon supplement them. Since information on post-translational modification is readily available from UniProt, some of these such as glycosylation and phosphorylation may be a very viable first target for modelling.

## Experimental

For our research we used the Google Colab^50^ “Colabfold: AlphaFold2 using MMseqs2” with the AlphaFold^7,51–53^ version v2.2.0 and non-premium access.

Clashscores, Ramachandran outliers and so-called “bad” bond lengths and angles were calculated with Molprobity^54^.

## Conclusions

The protein data bank represents the most important resource for structural biology, and machine-learning based fold prediction is now taking full advantage of this resource. In exchange, AlphaFold2 and more recently, RosettaFold^55^, can aid experimental design, facilitate structure solution and interpretation of maps, help to identify which part of a protein sequence may be intrinsically disordered. AlphaFold and RosettaFold are exciting new tools which will allow us to better understand macromolecular structures. Their shortcomings shine a spotlight on the shortcomings of our current experiments and how we interpret them with models. In order to push the boundaries of machine learning based fold prediction, we will need better training data. And this means that we need experiments and modelling methods that sample, for example, the entire conformational space of proteins. The methods themselves will also have to evolve, to include ligands, posttranslational modifications and complexes of different types of molecules. Nevertheless, machine learning based fold prediction are a game changer for structural bioinformatics and experimentalists alike, with exciting possibilities ahead.

## Author Contributions

Andrea Thorn: Supervision, Conceptualization, Formal analysis, Visualization, Investigation, Writing – Original draft, review & editing, Project administration, Funding acquisition. Dave Briggs: Writing – Original draft, review & editing. Max Edich: Methodology, Investigation, Formal analysis, Writing – editing. Oliver Kippes: Investigation, Writing - Original draft. Yunyun Gao: Data Curation & Resources, Writing - review and editing.

## Acknowledgements

The authors would like to thank Christopher Williams, Jane Richardson and Arwen Pearson for discussion. This work was supported by the German Federal Ministry of Education and Research (grant No. 05K19WWA) and the Deutsche Forschungsgemeinschaft (grant No. TH2135/21).

## Footnotes

‡ In the following we will only refer to AlphaFold 2 and its updates, although we write ‘AlphaFold’.

